# Over two orders of magnitude difference in rate of single chromosome loss among sundew (*Drosera* L., Droseraceae) lineages

**DOI:** 10.1101/2022.10.24.513289

**Authors:** Rebekah A. Mohn, Rosana Zenil-Ferguson, Thilo A. Krueger, Andreas S. Fleischmann, Adam T. Cross, Ya Yang

## Abstract

Chromosome number change is a driver of speciation in eukaryotic organisms. Carnivorous sundews, the plant genus *Drosera* L., exhibit single chromosome number variation among and within species, especially in the Australian *Drosera* subg. *Ergaleium* D.C., potentially linked to the presence of holocentromeres. We reviewed literature, verified chromosome counts, and using an *rbc*L chronogram, tested alternate models where the gain, loss, and doubling rates (+1, −1, ×2) were the same or different between *D*. subg. *Ergaleium* and the other subgenera. Ancestral chromosome number estimations were performed, and the distributions of self-compatibility and genome size were visualized across the genus. The best model for chromosome evolution had equal rates of polyploidy (0.014 per million years; Myr) but higher rates of single chromosome number gain (0.19 and 0.027 per Myr) and loss (0.23 and 0.00059 per Myr) in *D*. subg. *Ergaleium* compared to the other subgenera. We found no evidence for differences in single chromosome evolution to be due to differences in diploidization after polyploidy or to holocentromeres as had been proposed. This study highlights the complexity of factors influencing rates of chromosome number evolution.

---

Chromosome evolution events, such as duplication, inversion, fusion, and fission, are universal across the eukaryotic tree of life but appear to be more common in some lineages than others (reviewed in Coghlan et al., 2005). These chromosomal changes have long been considered driving forces of speciation and lineage diversification (Stebbins, 1971; Grant, 1981; Coyne and Orr, 2004). Therefore, identifying lineages with unusual rates of chromosome change and the intrinsic and environmental factors influencing these rates is critical to our understanding of evolutionary processes in general.

Recent developments in macroevolutionary modeling approaches have explored the association of chromosome evolution with trait evolution and lineage diversification (Mayrose et al., 2011; Freyman and Höhna, 2018; Baniaga et al., 2019; Zenil-Ferguson et al., 2019; Román-Palacios et al., 2020; Zhan et al., 2021). However, most of this work has focused on the role of chromosome doubling. Putative factors influencing the occurrence of single chromosome change include post-polyploidy dysploidy and rediploidization (Mandáková and Lysak, 2018), as well as centromere type (Luceño and Guerra, 1996; Mayrose and Lysak, 2020; Ruckman et al., 2020). Factors influencing the establishment of a new karyotype, such as autogamy (selfing) and clonality in plants, have only been explored in relation to polyploidy but likely impact single chromosome evolution as well (Husband et al., 2013; Weiss-Schneeweiss et al., 2013; Van Drunen and Husband, 2019). The relative importance of selfing and clonality in single chromosome evolution and establishment remains largely unknown.

Despite the importance of chromosome change to understanding evolution, obtaining a dataset of chromosome numbers with a matching phylogenetic tree to model the rates of chromosome change is challenging. A well-resolved phylogeny with a comprehensive species-level sampling is not always available. Further, because fresh root tips or flower buds are required to obtain chromosome counts, chromosome counts are often incomplete for lineages with wide geographic distributions. In addition to incomplete sampling, the quality of chromosome count datasets is eroded by chromosome counting errors (Windham and Yatskievych, 2003), reporting errors in chromosome number databases (Rivero et al., 2019), and taxonomic uncertainty from species misidentifications or taxonomic changes.

The carnivorous plants known as sundews (genus *Drosera* L.; family Droseraceae; order Caryophyllales) are exceptionally well-studied cytologically, with chromosome counts available for about half of its ca. 260 species. *Drosera* species are widely distributed and occur in a wide variety of habitats from boreal peatlands to tropical savannahs and subtropical sandplain heathlands and rock outcrops (Fleischmann et al., 2018). Hotspots of species diversity include Australia (ca. 170 species), Africa (ca. 40 species), and South America (ca. 40 species; Fleischmann et al., 2018). *Drosera* consists of four well-supported subgenera (Fleischmann et al., 2018): the two early-branching *D*. subg. *Regiae* Seine & Barthlott and *Arcturia* (Planch.) Schlauer harbor only one and two species respectively, while the sister *D*. subg. *Drosera* L. and *Ergaleium* D.C. are species-rich and harbor ca. 110 and ca. 150 species, respectively. Cytological studies on *Drosera* have been undertaken for over 120 years (Huie, 1897; Rosenberg, 1903), resulting in a rich literature record comprising more than 600 individual chromosome counts for ca. 140 species (e.g., Rothfels and Heimburger, 1968; Kress, 1970; Sheikh and Kondo, 1995; Chen, 1998; Rivadavia, 2005).

Previous karyotype studies in *Drosera* have revealed strikingly elevated levels of single chromosome number variation in *D*. subg. *Ergaleium* (almost every number from *n* = 3 to 23, with numbers up to 45; tuberous, pygmy, and woolly sundews of Australia; Table S1; Sheikh and Kondo 1995; Hoshi and Kondo, 1998; Rivadavia et al., 2003; Shirakawa, Hoshi, et al., 2011). In contrast, the other three subgenera exhibit primarily polyploid chromosome variation (Hoshi and Kondo, 1998; Rivadavia et al., 2003). The increased single chromosome number variation has been attributed to the presence of holocentric chromosomes in *Drosera* (Sheikh et al., 1995). Holocentric chromosomes have a centromere along their entire length rather than localized in the typical, X-shaped, monocentric chromosome. Because chromosomes of all *Drosera* except *D. regia* (Shirakawa, Nagano, et al., 2011) and *D. slackii* (Bennett and Cheek, 1990) lack a visible centromere constriction (Nontachaiyapoom et al., 2000; Kondo and Nontachaiyapoom, 2008), and all eight species tested so far undergo successful mitotic segregation after breakage (Sheikh et al., 1995; Furuta and Kondo, 1999; Shirakawa, Hoshi, et al., 2011; Zedek et al., 2016; Kolodin et al., 2018), researchers have hypothesized holocentromeres to be present in almost all *Drosera*. However, the distribution of phospho-histone 2A threonine-120, a histone commonly associated with the centromeric and pericentric region (Dong and Han, 2012; Wanner et al., 2015), indicates monocentromeres in three species from *D*. subg. *Drosera* and *D*. subg. *Ergaleium* (Demidov et al., 2014). Together, the evidence suggests that holocentromeres do not correspond to higher levels of chromosome number variation in *Drosera*. However, contrasting levels of chromosome number variation could also result from different ages of the lineages, uneven taxon sampling, counting errors, and taxonomic misidentification of material used for counts (e.g., the confusion of *D. aliciae* and *D. spatulata*; see Kress 1970; of *D. montana* and closely allied taxa; see Rivadavia, 2005). A critical evaluation of chromosome count data quality across all records is required to lay the foundations for subsequent analyses. Furthermore, the rate of chromosome number change has yet to be tested using a modeling framework that considers both the phylogenetic history and different modes of chromosome evolution. This phylogenetic modeling framework would also allow the investigation of associations between rates of chromosome number evolution and traits such as centromere type, life history, clonal propagation, and mating system.

In this study, we quantified the rate of chromosome doubling and single chromosome gain and loss on a dated phylogeny of *Drosera*. We tested whether the rates of chromosome evolution differ significantly between *D*. subg. *Ergaleium* and the other three subgenera, by critically evaluating previously published chromosome counts, verifying voucher specimens to identify possible taxonomic misidentifications, and using BiChrom (binary state linked to chromosome number change) models (Zenil-Ferguson et al., 2017) and Bayes factors to compare models of subgeneric differences in rates of chromosome evolution in a genus-wide phylogenetic context. An ancestral state reconstruction based on the resulting best-fit model was compared with genome size, life history, and centromere type to explore potential factors associated with different chromosome evolution rates between *Drosera* subgenera. Our analyses show highly elevated rates in single chromosome evolution but not polyploidy in *D*. subg. *Ergaleium* compared to the rest of the genus. Contrary to previous proposals, we found no evidence that such rate shift was due to diploidization after polyploidy or to holocentromeres, pointing to the complexity of factors contributing to rates of single chromosome evolution.

## METHODS

### Literature review and evaluation of chromosome counts

Lists of original references for *Drosera* chromosome counts were obtained from the Chromosome Counts Database (Rice et al., 2015), Index of Plant Chromosome Numbers (Goldblatt and Johnson, 1979–), citations referenced by additional publications on karyotypes in *Drosera* (Kondo, 1969; Dawson, 2000; Rivadavia et al., 2003; Veleba et al., 2017), and searches on Google Scholar and the library databases of the University of Minnesota, Curtin University, and University of Western Australia. Voucher specimen information, chromosome count methodology, and provenance data were recorded for every chromosome count identified either from the original publication or from subsequent literature in the case of 14 counts (six publications) where the original data could not be obtained.

Chromosome counts were excluded from analyses where the chromosome count methodology was flawed or original publication expressed uncertainty about the exact chromosome count (10 counts), where counts were made from primary hybrids (25 counts), or if there was taxonomic uncertainty about the material examined (73 counts). Taxonomic uncertainty was characterized by 1) counts that lack both species identification and voucher specimen; 2) species with taxonomy updates after the karyotype publication (especially in the case of species complexes), that lack sufficient provenance or character description and any voucher specimen with which to assign the taxon to the updated species name; 3) counts made from cultivated material of a species often misidentified in cultivation; or 4) a mismatch between the voucher specimen and the name associated with the count. See Supplemental Information S1 for details on evaluating published chromosome count data.

For species with two or more chromosome numbers after filtering, the number with the most counts was used for subsequent modeling analyses. For 11 species where multiple chromosome numbers had an equal number of counts, one value was selected at random.

### Phylogenetic reconstruction for comparative analyses

In order to estimate a chronogram for modeling chromosome number evolution, *rbc*L sequences for *Drosera* species and outgroup taxa for non-core Caryophyllales were retrieved from the GenBank (Table S2). Five sequences were removed due to ambiguous nucleotide sites. The taxonomy for *rbc*L sequences with herbarium vouchers at M and SPF (herbarium acronyms following Index Herbariorum) were updated as noted in Table S2. For species with multiple *rbc*L sequences, the longest sequence was kept.

Sequences were aligned with default settings using the MAFFT (Katoh and Standley, 2013) plug-in for Geneious version 11.1.5 (Kearse et al., 2012). The ends of sequences that were only present in two outgroup species were trimmed. Priors for molecular dating in BEAST version 2.6.4 (Bouckaert et al., 2014) followed previous molecular dating analysis across the Caryophyllales (Yao et al., 2019) using a lognormal relaxed molecular clock and the birth-death model of speciation. For each fossil, the prior node was constrained to a lognormal distribution with a mean of 1.0, a standard deviation of 0.5, and an offset based on the age of the fossil. As in Yao et al. (2019), fossil *Aldrovanda intermedia* and *A. ovata* was used to set the prior for the most recent common ancestor (MRCA) of *Dionaea* and *Aldrovanda* with an offset of 41.2 Ma, and *Polygonocarpum johnsonii* was used to constrain the MRCA of the Polygonoideae (in Polygonaceae) included with an offset of 66.0 Ma. The MRCA of non-core Caryophyllales was constrained to 115 Ma with a normal distribution and a standard deviation of 4.0 Ma representing the 95% confidence interval in the posterior distribution of the dating analysis of Yao et al. (2019). The Markov-Chain Monte Carlo (MCMC) was run for 100,000,000 generations, sampling every 1000 generations. The BEAST input file and data are available at 10.5281/zenodo.6081366. The resulting summary statistics were visualized in Tracer version 1.7.1 (Rambaut et al., 2018). The obtained phylogenetic trees were further reduced to 1 in 10 and summarized in TreeAnnotator version 2.6.2 (Drummond and Rambaut, 2007) with a 10% burn-in, and the maximum clade credibility tree was visualized in FigTree version 1.4.4 (Rambaut, 2018). The chronogram (using the ape R package; Paradis and Schliep, 2019) and chromosome count matrices were trimmed to species shared by both datasets for subsequent analyses.

### Modeling chromosome number evolution

We used the binary trait linked to chromosome number change model (BiChrom; Zenil-Ferguson et al. 2017) and implemented it in RevBayes software version 1.1.0 (Höhna et al., 2016) to estimate the differences in three rates of chromosome number evolution for each binary state (Fig. 1): γ (a single chromosome gain, by duplication or fission), δ (a single chromosome loss, by rearrangement, fusion, or loss), and ρ (a polyploidy event). The binary state is defined as whether a taxon belongs to *D*. subg. *Ergaleium* (state E) or not, in which case it belongs to *D*. subg. *Drosera*, *Arcturia*, or *Regiae* (state D). By defining our binary state in this fashion, we estimate a transition rate q, which is a nuisance parameter but allows us to correctly compare rates of chromosome change between the two groups. Species were assigned as state E or state D sensu Fleischmann et al. (2018).

**Fig. 1:**
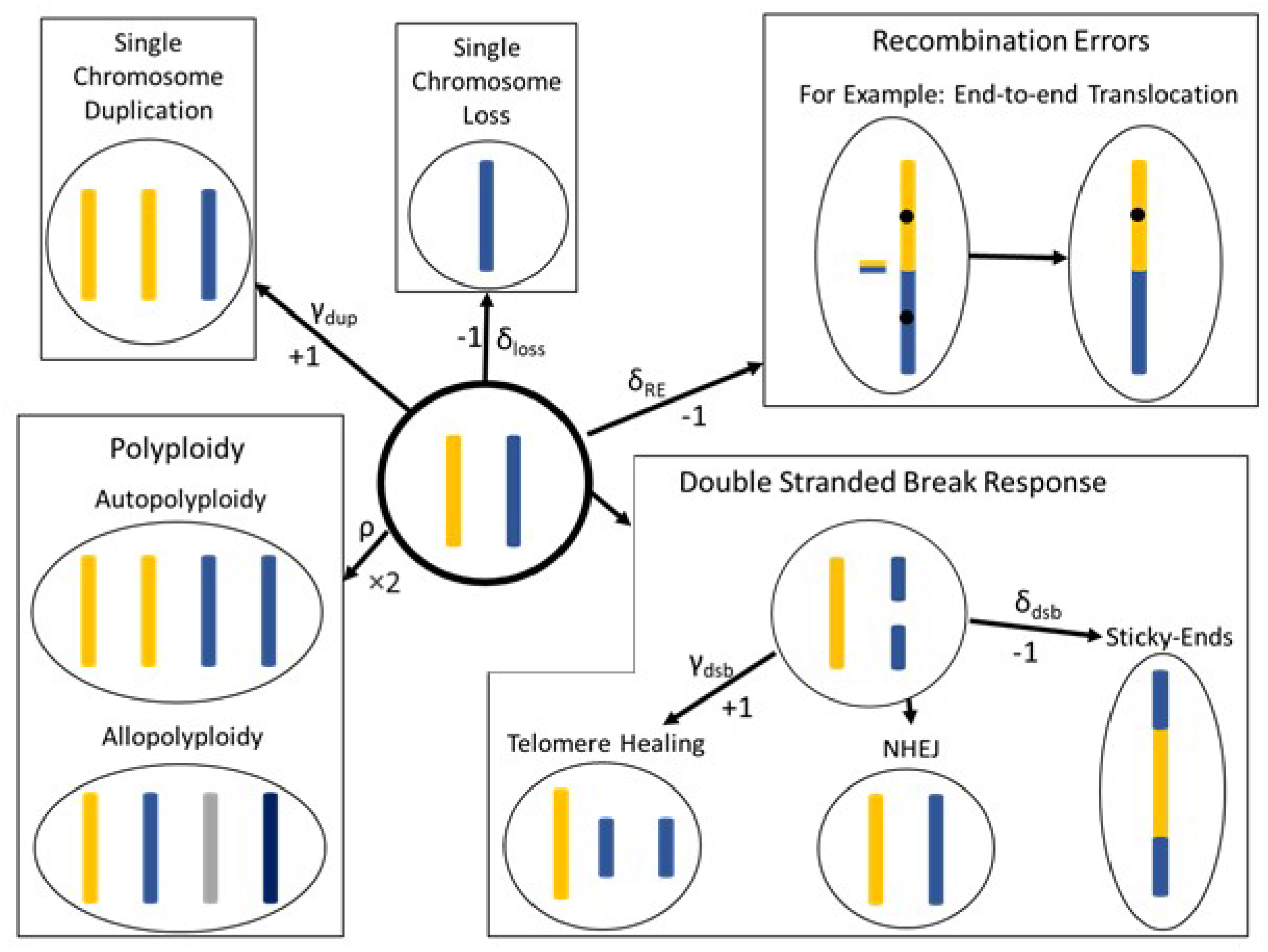
A summary of the processes that can give rise to changes in chromosome number. Each cell is depicted in haploid form. The original cell (center) starts with two haploid chromosomes. Arrows indicate changes in chromosomes and, where possible, are labelled with the type of change (+1, −1, ×2) and the symbol used in BiChrom models (γ, δ, and ρ respectively; Mayrose & Lysak, 2020). The centromere is shown as a black spot in the recombination error to emphasize the steps required to handle an additional centromere. Since +1 and −1 can occur via multiple mechanisms with different impacts on gene copy number, for example, a subscript is used to distinguish the cause of change. An increase in one chromosome can be due to telomere healing after a chromosome break or a single chromosome duplication; a single chromosome decrease can be due to a recombination error (Nested insertion, end-to-end translocation, or Robertsonian Translocation; Mayrose & Lysak, 2020), two chromosomes fusing after a breakage, or the loss of a single chromosome. Single chromosome loss is unlikely except after polyploidy (Luceno & Guerrra, 1996). A doubling of all chromosomes can be due to an auto- or allo-polyploidy. Holocentromeres are expected to alleviate issues caused by acentric fragments after double stranded breaks and tangling of bicentric chromosomes after fusion (Cuacos et al., 2015).

Our macroevolutionary modeling framework involved simultaneously estimating the rate of evolution of chromosome number and a binary state along a phylogeny. We first defined a matrix describing the instantaneous rate of chromosome number change between two chromosome numbers and between the two states at the same chromosome number (Fig. S1; Mayrose et al., 2010; Zenil-Ferguson et al., 2017). Commonly known as the Q-matrix for continuous time Markov chains, this matrix can be numerically difficult to use because chromosome transition matrices are large and contain many zeros since transitions reflect only single chromosome number changes or doubling (Mayrose et al., 2010; Zenil-Ferguson et al., 2017). These types of matrices are numerically unstable when exponentiated, so limiting the maximum number of chromosomes and rates included is key for estimation (Zenil-Ferguson et al., 2018). Therefore, in our dataset with chromosome number (2*n*) ranging from 8 to 60, we first calculated haploid chromosomes (1*n*) and set the chromosome states for the Q-matrix ranging from 1 to 35 and a bin for 35+ haploid chromosomes to make the matrix more computationally stable (Fig. S1; Zenil-Ferguson et al., 2017, Zenil-Ferguson et al., 2018). We removed *Drosera lanata* (2*n* = 19), to avoid non-integer haploid chromosome numbers, and records of B-chromosomes, as these small satellite chromosomes do not segregate normally during cell division. The resulting matrix had 72 rows and 72 columns reflecting 1 to 35 and 35+ chromosome numbers for both states E and D (Fig. S1). Since we expect the chromosome evolution rate in *Drosera* outside of *D*. subg. *Ergaleium* to be more similar to the rate in most angiosperms, we considered state D the ancestral state and E the derived state and only allowed transitions from state D to state E. The probabilities of the root being 1 to 35+ chromosomes in either state were set equal.

Three nested models were used for comparison, each with a subset of the rates being constrained as equal across the two states. The full model (H2) allowed rates (ρ = chromosome doubling, δ = chromosome loss, γ = chromosome gain) to vary as a function of each of the states D or E. The fixed-polyploid model (H1; ρ_D_ = ρ_E_) constrained the rate of chromosome doubling to be the same in D and E. The null model (H0) constrained all rates to be equal for the two states (ρ_D_ = ρ_E,_ γ_D_ = γ_E,_ δ_D_ = δ_E_). Rate priors for all chromosome transitions in both states were set to an exponential distribution with a mean of 1/3 probability of change per million years (Myr).

We ran our custom MCMC scripts in RevBayes (Höhna et al., 2016) for more than 200,000 generations until convergence was reached and checked using Tracer (Rambaut et al., 2018). We also verified that effective sample sizes for all the parameters were above 200. Concurrently, we reconstructed ancestral states using marginal posterior probabilities for each of the internal nodes as part of the inference following Freyman and Höhna (2018) and Zenil-Ferguson et al. (2019). The RevBayes input data and scripts are at 10.5281/zenodo.6081366.

The three models were compared using Bayes factors in RevBayes (Höhna et al., 2016) by calculating power posterior distributions with twenty stepping-stones (Xie et al., 2011). The stepping-stone algorithm was used to calculate the marginal likelihood of each model by estimating the probability of the data between the prior and the posterior. This is done by raising the posterior distribution of the MCMC to a power ranging from 0 to 1, thus providing a discrete approximation between the prior and posterior probabilities. The marginal log likelihoods were calculated from these stepping-stones and were then subtracted to calculate the Bayes factors *κ* statistic. *κ* > 6 is strong evidence in favor of the model input first in the calculation of x is assumed. If *κ* > 1, there is moderate support, and no evidence in favor of either model if *κ* is between −1 and 1 as described in Kass and Raftery (1995). If *κ* results in large negative values, the evidence goes in favor of the model whose marginal log-likelihood is subtracting in the calculation of *κ*.

All the MCMC outputs were analyzed using Tracer with a burn-in of 10% discarded. The resulting ancestral state reconstruction for the best supported model was visualized with RevGadgets R package (Tribble et al., 2021).

### Genome size and mating system

Self-compatibility data for 98 species of *Drosera* were obtained from publications (Table S3). Recent studies (Fleischmann, in press) suggest all *D. auriculata* are self-compatible contrary to a (doubtful) previously-published report by Chen et al. (1997). *Drosera* genome sizes were obtained from Veleba et al. (2017), or newly generated in this study for 17 species at the Flow Cytometry Core Lab at the Benaroya Research Institute (Seattle, WA, U.S.A.). Source, voucher, and size standards used for generating new flow cytometry data are listed in Table S3.

## RESULTS

### *Chromosome Counts for 127* Drosera *species show distinctive patterns of variation between* D. *subgenus* Ergaleium *and other subgenera*

An initial dataset of 676 chromosome counts in *Drosera* from 150 species or hybrids were compiled (Table S1). After removing hybrids and low-quality counts, 510 counts from 127 species were used for downstream analyses (ca. 48% of all species). These counts included 32% of named species from Africa, 45% from South America, 51% from Australia, 60% from Asia, and all species from North America and Europe. *Drosera* subg. *Arcturia*, *Drosera*, *Ergaleium*, and *Regiae* had respectively 50%, 43%, 51%, and 100% of species with counts.

Almost every even chromosome number from 2*n* = 6 to 46 was reported from *D*. subg. *Ergaleium*, including within-species variation. In contrast, *D*. subg. *Drosera* has chromosome numbers from 2*n* = 16 to 80 with variation primarily in polyploid series (2*n* = 20, 40, 60, 80; Fig. 2). Chromosome number for *D. arcturi* (*D. subg. Arcturia*) was 2*n* = 20 and for *D. regia* (*D*. subg. *Regiae*) was 2*n* = 34 (Fig. 2; Table S1).

**Fig. 2:**
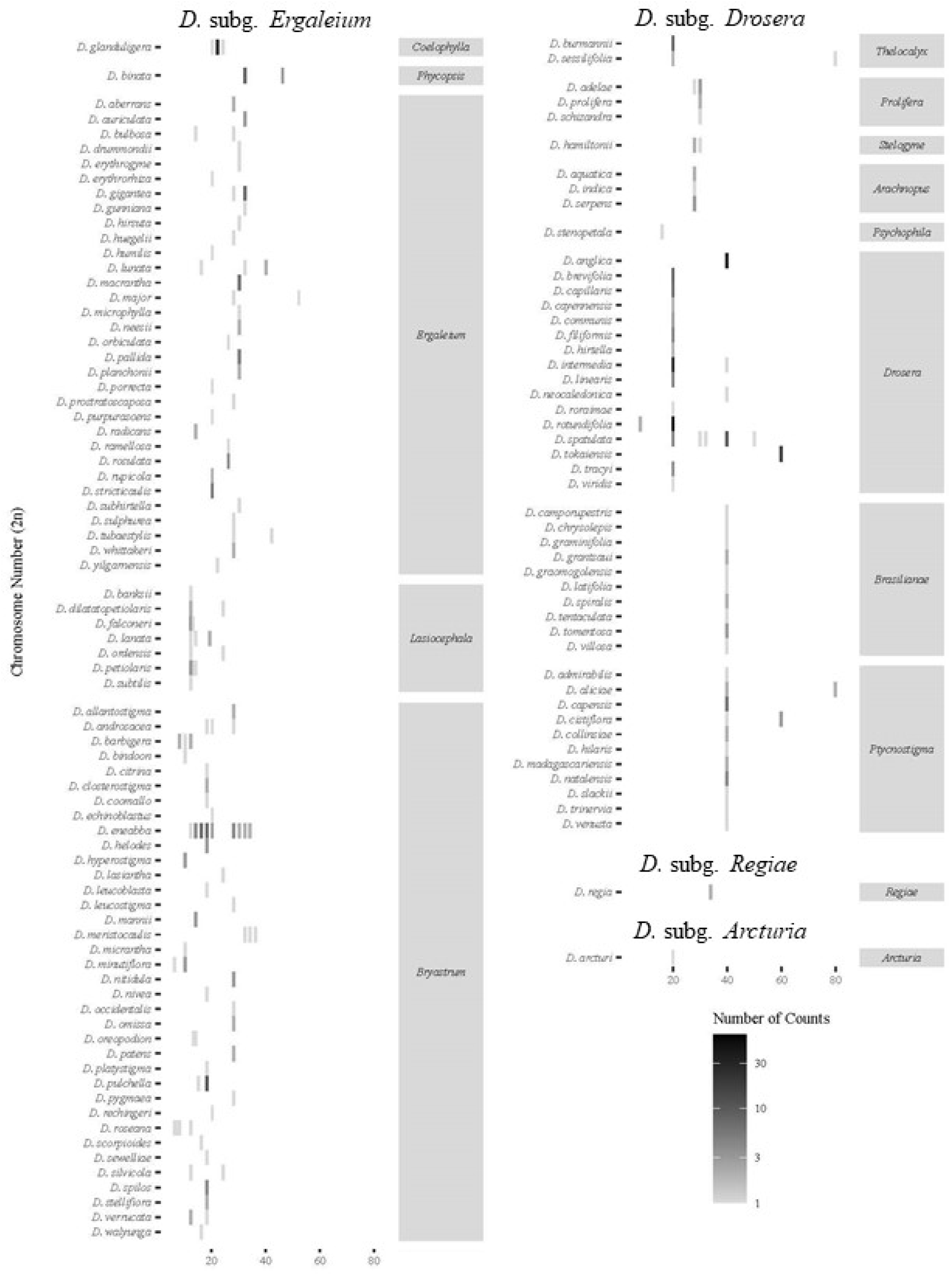
*Drosera* subg. *Ergaleium* (left) exhibits marked chromosome variation within sections, and even within species. Little within species or within section variation was observed for *D*. subg. *Drosera* (right), and where variation was observed it fell primarily into polyploidy series. The shade of the square indicates the number of samples for each species, emphasizing that the lack of variation within *D*. subg. *Drosera* is not due to a lack of samples.

### Chronogram Reconstruction

The trimmed *rbc*L matrix included 1,440 bases with 478 variable sites across the 17 outgroup and 79 ingroup taxa. After burn-in, the ESS in BEAST was greater than 200 for all continuous statistics. The *rbc*L tree placed *D. regia* and *D. arcturi* in a clade with *Aldrovanda* and *Dionaea* that was sister to the rest of *Drosera* with strong to moderate support, likely due to long-branch attraction. BEAST analysis estimated the crown age of *D*. subg. *Ergaleium* at 52.0 Mya and *D*. subg. *Drosera* at 49.6 Mya (Fig. S2).

### Drosera *subgenus* Ergaleium *differs from other subgenera in chromosome evolution rate*

We had available chromosome counts and phylogenetic *rbc*L data for 59 species: 25 species from *D*. subg. *Ergaleium*, 32 species from *D*. subg. *Drosera*, and one species each from *D*. subg. *Arcturia* and *D*. subg. *Regiae*. The BiChrom analysis for the full model with all rates estimated separately between *D*. subg. *Ergaleium* and the other subgenera (H2) took over 200,000 generations to converge as the posterior distribution was bimodal.

In the full model (H2), the mean of the posterior probabilities of gaining (γ_E_ = 0.23) or losing (δ_E_ = 0.25) one chromosome in *D*. subg. *Ergaleium* was 8.8-fold and 40.3-fold higher than other subgenera (γ_D_ = 0.026; δ_D_ = 0.0062; Table S4; Fig. 3). These rates are interpreted as the amount of single chromosome change per million years. However, the rate of chromosome gain for *D*. subg. *Drosera*, *Arcturia*, and *Regiae* falls within the first quartile of the rate of chromosome gain for *D*. subg. *Ergaleium* and only the 95% credible interval for the rates of single chromosome loss was distinct (95% HPD δ_E_ = 0.063 to 0.52; 95% HPD δ_D_ = 6.2×10^−6^ to 4.4×10^−2^; Table S4; Fig. 3). The rates of polyploidy largely overlapped (Fig. 3).

**Fig. 3:**
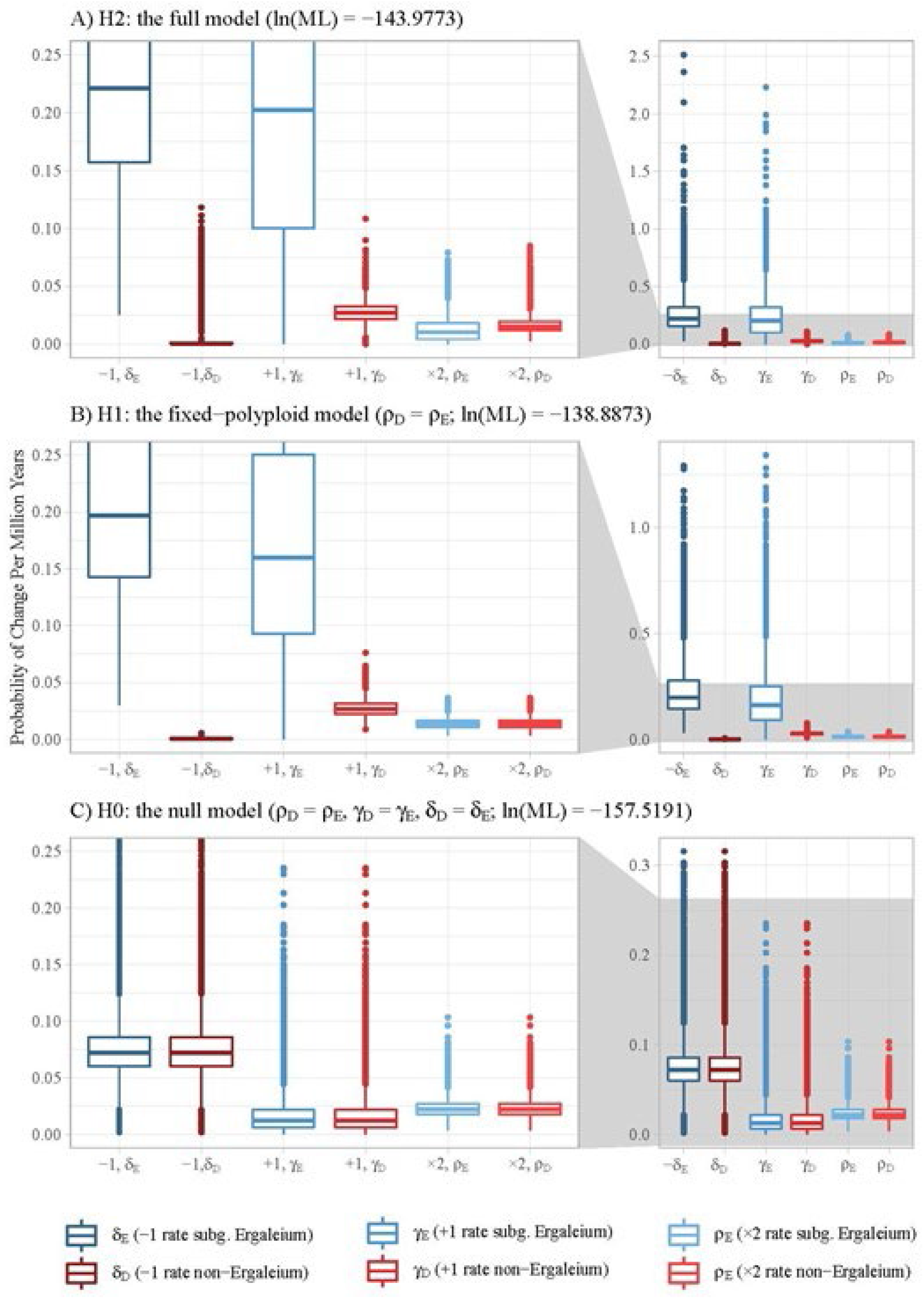
The posterior sampling distribution from the MCMC after burn-in for (A) H2 where all rates were estimated independently for *Drosera* subg. *Ergaleium* (state E) from the other three *Drosera* subgenera (state D), (B) H1 where all rates were independent except ρ (polypidy) which equal across *Drosera* and (C) H0 where all rates were equal across *Drosera*. δ_D_ and δ_E_ are significantly distinct in H2 and H1. All remaining rates were not significantly different between state E and D.

Compared to rates estimated in the full model, the null model (H0) estimated an intermediate rate for losing one chromosome, while the estimated rate of polyploidy doubled and the rate for gaining a chromosome decreased (Fig. 3). Comparing Bayes factors for the full model and null model found that the full model had strong support (BF =13.5), showing supporting evidence that there are differences between *D*. subg. *Ergaleium* and the other subgenera.

### *Rate of polyploidy does not differ among* Drosera *subgenera*

Given the very similar inferred chromosome doubling rates for all the subgenera and the genomic instability and potential chromosome loss post a polyploidy event, we tested an additional model estimating chromosome loss and gain for the two groups separately but polyploidy together (H1). The MCMC run for H1 had an effective sampling size above 200 for all statistics and solved issues with the bimodality found in model H2. We found a moderate preference for H1 over the full model (H2; BF = 5.1; Table S4; Fig. 3).

The best fit model (H1) with both chromosome loss (δ) and gain (γ) as functions of each of the subgenera showed higher chromosome loss and chromosome gain rates in *D*. subg. *Ergaleium* and 95% credible intervals similar to the full model (Table S4; Fig. 3). The mean δ_E_ was 389.8-fold higher than δ_D_ and the 95% HPD did not overlap (Table S4; Fig. 3). With overlapping 95% HPDs, the mean γ_E_ was over 6-fold different than γ_D_ (Table S4; Fig. 3).

Under the H1 model, the ancestral state reconstruction estimated the MRCA of *Drosera* to have a haploid chromosome number of eight and state D. The base of *D*. subg. *Ergaleium* also had a haploid chromosome number of eight but with state E. The difference in single chromosome change between subgenera is supported across the reconstruction by the stability of chromosome number in *D*. subg. *Drosera* and repeated changes in *D*. subg. *Ergaleium*. Based on the reconstruction, polyploidization events occurred five times in *D*. subg. *Ergaleium*, three times in *D*. subg. *Drosera*, and once in *D*. subg. *Regiae* (Fig. 4).

**Fig. 4:**
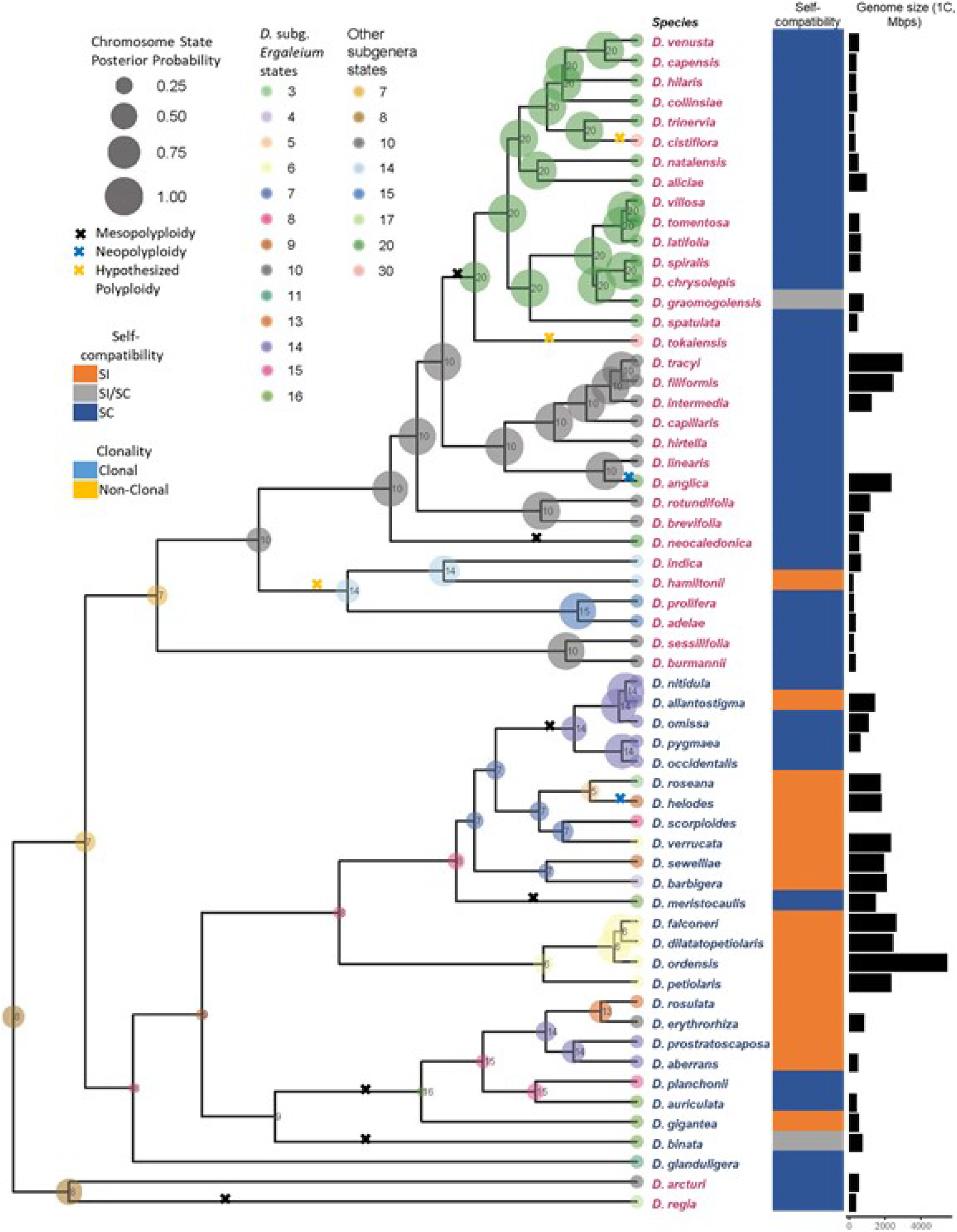
In addition to having higher rates of single chromosome change, *Drosera* subg. *Ergaleium* (species name in blue) have more species that reproduce clonally and are self-incompatible than the other three subgenera (species name in red). Most polyploid lineages have smaller genome sizes than their diploid sister lineages (marked with a black × on tree). Numbers and colored bubbles on nodes and tips were ancestral state reconstruction of the chromosome numbers and binary state. Size of bubbles indicate the posterior probability of number. Clonality (occurring in a section with a structure for reproducing genetically identical individuals), self-incompatibility (the ability or inability to produce viable offspring when crossed with itself), and diploid genome size were presented to the right of the species names. Polyploidy as seen by chromosome complete doubling (older and smaller genomes than sister lineages: black ×; recent and equal too or bigger than sister lineages: blue ×) and 1.5 duplication (yellow ×) were marked on branches.

### *Self-compatibility differs between* Drosera *subgenera*

In *D*. subg. *Ergaleium*, 48 of the 60 (80%) species with known mating systems are self-incompatible in at least some populations (Fig. 4; Table S3.2). In contrast, only three of the 38 species (8%) in the remaining three subgenera are self-incompatible, both of them are in *D*. subg. *Drosera* but not closely related (Fig. 4; Table S3.2).

## DISCUSSION

### *Rates of single chromosome number change significantly differ among* Drosera *subgenera*

After correcting for counting and taxonomic errors and using a model that considers time, the rate of polyploidy in *Drosera* (0.014 per Myr) did not differ between subgenera and was very similar to the polyploidy rate previously reported for perennial angiosperms (0.015 per Myr; Van Drunen and Husband, 2019) and median rate across angiosperm families (0.025 per Myr; Zhan et al., 2021). Similarly, the single chromosome gain (0.027) and loss rate (0.00059) for *Drosera* lineages except *D*. subg. *Ergaleium* fell higher and lower, respectively, than the average rate (0.0061 and 0.016 respectively) across angiosperm families (Zhan et al., 2021). In contrast, the rate of single chromosome shifts in *D*. subg. *Ergaleium* was 6-fold (chromosome gain) and 350-fold (chromosome loss) higher than in the remainder of the genus, and the rates of *D*. subg. *Ergaleium* are likely even higher with increased species sampling. Orders of magnitude differences in chromosome loss and gain rates have also been documented between herbaceous versus woody plants, among some *Carex* lineages and among some insect lineages (Escudero et al., 2014; Zenil-Ferguson et al., 2017; Ruckman et al., 2020; Sylvester et al., 2020).

Elevated rates of single chromosome evolution can be due to increased rates of polyploidy and subsequent rediploidization (Mandáková and Lysak, 2018). However, we found no difference in rates of polyploidy among subgenera in *Drosera*. Although polyploid species in *D*. subg. *Drosera* were considered stable polyploids as their chromosome numbers follow polyploid series (Hoshi and Kondo, 1998; Shirakawa, Hoshi, et al., 2011), we found evidence for genome downsizing after polyploidy across *Drosera*. Of the nine polyploidy events inferred, the most recent (ca. 3.3 Mya) has a genome size close to double that of the sister lineage, while the remaining eight more ancient polyploid lineages have similar or, in seven cases, smaller genome sizes than their diploid sister lineages (Fig. 4; Table S3.1; Veleba et al., 2017). Therefore, both the rate of polyploidy and the post-polyploidy diploidization show similar patterns across *Drosera* and no evidence supports either being the major cause in single chromosome number shifts in *Drosera*.

While the chromosome loss rates were clearly distinct between *D*. subg. *Ergaleium* and the other subgenera, the credible intervals for chromosome gain rates overlapped. Our analysis included 23% of the named species. To narrow the credible intervals, it is critical for future molecular and cytological work to include proper vouchers, locality information, the number of cells, and individuals counted.

### Towards drivers of chromosome evolution rate shift

Holocentromeres have been associated with increased chromosome fission producing a higher number of smaller chromosomes (Cuacos et al., 2015; Ruckman et al., 2020) as chromosome fragments with centromeres can pair and segregate properly even in heterozygous individuals (Luceño and Guerra, 1996; Jankowska et al., 2015; Ruckman et al., 2020). So far, no experimental evidence supports *D*. subg. *Ergaleium* having a distinct centromere type from the rest of the genus. This lack of association between holocentromeres and significant differences in chromosome evolution rates was also documented in insects (Ruckman et al., 2020).

A newly formed karyotype may be eliminated due to drift or selection against the deleterious nature of heterozygous individuals, especially in monocentric plants (Husband et al., 2013). Species with means of reproductive assurances (clonal propagation, selfing, etc.) may avoid these issues as the proportion of individuals in the population with the new chromosome number can increase without producing heterozygous individuals (Husband et al., 2013; Van Drunen and Husband, 2019; Spoelhof, Keeffe, et al., 2020). While a perennial life history and clonal propagation are common across *Drosera* (Fleischmann et al. 2018), contrary to expectation, a higher percentage of species studied in *D*. subg. *Ergaleium* are self-incompatible compared to the other subgenera (Fig 4; Table S3). Interestingly, Spoelhof, Keeffe, et al. (2020) proposed that sexual reproduction (especially outcrossing) is important for the long-term maintenance of species diversity after the formation of a new karyotype.

Moving forward, exploring the factors that are typically considered within a single species, such as population size, spatial distribution, and meiotic drive, would help dissect the mechanisms underlie new karyotype establishment and macroevolutionary diversification in *Drosera* and beyond (Reed et al., 2013; Bureš and Zedek, 2014; Blackmon et al., 2019; Ruckman et al., 2020; Spoelhof, Soltis, et al., 2020; Griswold, 2021).

### Conclusion

Differences in chromosome number variation between *Drosera* subg. *Ergaleium* and *D*. subg. *Drosera, Arcturia*, and *Regiae* result from significant differences in single chromosome evolution rate rather than sampling bias, chromosome counting errors, or clade age. *D*. subg. *Ergaleium* not only exhibits highly accelerated single chromosome evolution but also a higher percentage of self-incompatible species. Future work on both the natural history and molecular fronts are needed to tease apart the mechanisms underlying the highly elevated rate of single chromosome change. More broadly, our findings illustrate that additional factors other than genome downsizing after polyploidy and holocentromeres impact the rate of single chromosome evolution.

## Acknowledgements

The authors thank Fernando Rivadavia for his voucher identification and feedback on the manuscript, Alex Eilts and the College of Biological Sciences Conservatory for their assistance in growing plants for genome size estimation, Aaron Lee for his feedback on the manuscript, the University of Western Australia library for access to Lin Chen’s Thesis, Sergey Matveev for help with Russian translation, and Virginia’s Department of Conservation and Recreation and Darren Loomis for access and help obtaining specimens to the Cherry Orchard Bog Natural Area Preserve. The work is supported by the National Science Foundation (Grant Number: 2015210), the Fulbright Futures Program, and the Botanical Society of America.

## SUPPLEMENTAL MATERIALS

Figure S1: The transition matrix. See Fig. 1 for definition of chromosome transition parameters. Q_10_ is the transition state from state 1 to state 0.

Figure S2: The dated *rbc*L phylogeny with all taxa included from the BEAST analysis. Bars on nodes represent the 95% HPD intervals for the age of the node.

Table S1: The chromosome count data matrix with notes. Table S1.1 is the matrix itself, Table S1.2 contains the headers and information, and Table S1.3 contains the references for all the data.

Table S2: Source for *rbc*L sequences (Table S2.1) including the species name used, the GenBank ID, the originally reported species name, and the reason for taxonomic change if applicable. The species authority for each *Drosera* species (Table S2.2)

Table S3: The genome size (Table S3.1), self-compatibility (Table S3.2), and reference (Table S3.3). The genome size matrix included species names, locality and voucher information (visit 10.5281/zenodo.6081366 for photo vouchers), control, and reference. The self-compatibility data included species, reference, and notes on changes in taxonomy.

Table S4: Table S4.1 shows the marginal log likelihood; estimated rate of chromosome loss (δ), gain (γ), and polyploidy (ρ) for both state E (*Drosera* subg. *Ergaleium*) and state D (the other three *Drosera* subgenera); and the transition from state D to state E for three models. In the full model (H2) all the rates are estimated independently. Table S4.2 contains the 95% HPD distributions for the three models.

Supplemental Information S1: Methods for the chromosome count scoring and filtering.

## Notes

Conflict of Interests: The authors have no conflict of interests to declare.

DOI:10.5281/zenodo.6081366

### Competing Interest Statement

The authors have declared no competing interest.

