## Supplemental Figure 1 for "Over two orders of magnitude difference in rate of single chromosome loss among sundew (*Drosera* L., Droseraceae) lineages"

| | | | State E | | | | | | | | | | | | | | | | | | | | | | | | | | | | | | | | | | | |
|---|---|---|---|---|---|---|---|---|---|---|---|---|---|---|---|---|---|---|---|---|---|---|---|---|---|---|---|---|---|---|---|---|---|---|---|---|---|---|
| | | n | 1 | 2 | 3 | 4 | 5 | 6 | 7 | 8 | 9 | 10 | 11 | 12 | 13 | 14 | 15 | 16 | 17 | 18 | 19 | 20 | 21 | 22 | 23 | 24 | 25 | 26 | 27 | 28 | 29 | 30 | 31 | 32 | 33 | 34 | 35 | 35+ |
| | n | | 1 | 2 | 3 | 4 | 5 | 6 | 7 | 8 | 9 | 10 | 11 | 12 | 13 | 14 | 15 | 16 | 17 | 18 | 19 | 20 | 21 | 22 | 23 | 24 | 25 | 26 | 27 | 28 | 29 | 30 | 31 | 32 | 33 | 34 | 35 | 36 |
| State E | 1 | 1 ||$\gamma_0+\rho_0$|||||||||||||||||||||||||||||||||||
| | 2 | 2 |$\delta_0$||$\gamma_0$|$\rho_0$||||||||||||||||||||||||||||||||
| | 3 | 3 ||$\delta_0$||$\gamma_0$||$\rho_0$||||||||||||||||||||||||||||||
| | 4 | 4 |||$\delta_0$||$\gamma_0$|||$\rho_0$||||||||||||||||||||||||||||
| | 5 | 5 ||||$\delta_0$||$\gamma_0$||||$\rho_0$||||||||||||||||||||||||||
| | 6 | 6 |||||$\delta_0$||$\gamma_0$|||||$\rho_0$||||||||||||||||||||||||
| | 7 | 7 ||||||$\delta_0$||$\gamma_0$||||||$\rho_0$||||||||||||||||||||||
| | 8 | 8 |||||||$\delta_0$||$\gamma_0$|||||||$\rho_0$||||||||||||||||||||
| | 9 | 9 ||||||||$\delta_0$||$\gamma_0$||||||||$\rho_0$||||||||||||||||||
| | 10 | 10 |||||||||$\delta_0$||$\gamma_0$|||||||||$\rho_0$||||||||||||||||
| | 11 | 11 ||||||||||$\delta_0$||$\gamma_0$||||||||||$\rho_0$||||||||||||||
| | 12 | 12 |||||||||||$\delta_0$||$\gamma_0$|||||||||||$\rho_0$||||||||||||
| | 13 | 13 ||||||||||||$\delta_0$||$\gamma_0$||||||||||||$\rho_0$||||||||||
| | 14 | 14 |||||||||||||$\delta_0$||$\gamma_0$|||||||||||||$\rho_0$||||||||
| | 15 | 15 ||||||||||||||$\delta_0$||$\gamma_0$||||||||||||||$\rho_0$||||||
| | 16 | 16 |||||||||||||||$\delta_0$||$\gamma_0$|||||||||||||||$\rho_0$||||
| | 17 | 17 ||||||||||||||||$\delta_0$||$\gamma_0$||||||||||||||||$\rho_0$||
| | 18 | 18 |||||||||||||||||$\delta_0$||$\gamma_0$||||||||||||||||$\rho_0$|
| | 19 | 19 ||||||||||||||||||$\delta_0$||$\gamma_0$|||||||||||||||$\rho_0$|
| | 20 | 20 |||||||||||||||||||$\delta_0$||$\gamma_0$||||||||||||||$\rho_0$|
| | 21 | 21 ||||||||||||||||||||$\delta_0$||$\gamma_0$|||||||||||||$\rho_0$|
| | 22 | 22 |||||||||||||||||||||$\delta_0$||$\gamma_0$||||||||||||$\rho_0$|
| | 23 | 23 ||||||||||||||||||||||$\delta_0$||$\gamma_0$|||||||||||$\rho_0$|
| | 24 | 24 |||||||||||||||||||||||$\delta_0$||$\gamma_0$||||||||||$\rho_0$|
| | 25 | 25 ||||||||||||||||||||||||$\delta_0$||$\gamma_0$|||||||||$\rho_0$|
| | 26 | 26 |||||||||||||||||||||||||$\delta_0$||$\gamma_0$||||||||$\rho_0$|
| | 27 | 27 ||||||||||||||||||||||||||$\delta_0$||$\gamma_0$|||||||$\rho_0$|
| | 28 | 28 |||||||||||||||||||||||||||$\delta_0$||$\gamma_0$||||||$\rho_0$|
| | 29 | 29 ||||||||||||||||||||||||||||$\delta_0$||$\gamma_0$|||||$\rho_0$|
| | 30 | 30 |||||||||||||||||||||||||||||$\delta_0$||$\gamma_0$||||$\rho_0$|
| | 31 | 31 ||||||||||||||||||||||||||||||$\delta_0$||$\gamma_0$|||$\rho_0$|
| | 32 | 32 |||||||||||||||||||||||||||||||$\delta_0$||$\gamma_0$||$\rho_0$|
| | 33 | 33 ||||||||||||||||||||||||||||||||$\delta_0$||$\gamma_0$|$\rho_0$|
| | 34 | 34 |||||||||||||||||||||||||||||||||$\delta_0$||$\gamma_0$|$\rho_0$|
| | 35 | 35 ||||||||||||||||||||||||||||||||||$\delta_0$||$\gamma_0+\rho_0$|
| | 35+ | 36 |||||||||||||||||||||||||||||||||||$\delta_0$||
| State D | 1 | 37 |$q_{10}$|||||||||||||||||||||||||||||||||||
| | 2 | 38 ||$q_{10}$||||||||||||||||||||||||||||||||||
| | 3 | 39 |||$q_{10}$|||||||||||||||||||||||||||||||||
| | 4 | 40 ||||$q_{10}$||||||||||||||||||||||||||||||||
| | 5 | 41 |||||$q_{10}$|||||||||||||||||||||||||||||||
| | 6 | 42 ||||||$q_{10}$||||||||||||||||||||||||||||||
| | 7 | 43 |||||||$q_{10}$|||||||||||||||||||||||||||||
| | 8 | 44 ||||||||$q_{10}$||||||||||||||||||||||||||||
| | 9 | 45 |||||||||$q_{10}$|||||||||||||||||||||||||||
| | 10 | 46 ||||||||||$q_{10}$||||||||||||||||||||||||||
| | 11 | 47 |||||||||||$q_{10}$|||||||||||||||||||||||||
| | 12 | 48 ||||||||||||$q_{10}$||||||||||||||||||||||||
| | 13 | 49 |||||||||||||$q_{10}$|||||||||||||||||||||||
| | 14 | 50 ||||||||||||||$q_{10}$||||||||||||||||||||||
| | 15 | 51 |||||||||||||||$q_{10}$|||||||||||||||||||||
| | 16 | 52 ||||||||||||||||$q_{10}$||||||||||||||||||||
| | 17 | 53 |||||||||||||||||$q_{10}$|||||||||||||||||||
| | 18 | 54 ||||||||||||||||||$q_{10}$||||||||||||||||||
| | 19 | 55 |||||||||||||||||||$q_{10}$|||||||||||||||||
| | 20 | 56 ||||||||||||||||||||$q_{10}$||||||||||||||||
| | 21 | 57 |||||||||||||||||||||$q_{10}$|||||||||||||||
| | 22 | 58 ||||||||||||||||||||||$q_{10}$||||||||||||||
| | 23 | 59 |||||||||||||||||||||||$q_{10}$|||||||||||||
| | 24 | 60 ||||||||||||||||||||||||$q_{10}$||||||||||||
| | 25 | 61 |||||||||||||||||||||||||$q_{10}$|||||||||||
| | 26 | 62 ||||||||||||||||||||||||||$q_{10}$||||||||||
| | 27 | 63 |||||||||||||||||||||||||||$q_{10}$|||||||||
| | 28 | 64 ||||||||||||||||||||||||||||$q_{10}$||||||||
| | 29 | 65 |||||||||||||||||||||||||||||$q_{10}$|||||||
| | 30 | 66 ||||||||||||||||||||||||||||||$q_{10}$||||||
| | 31 | 67 |||||||||||||||||||||||||||||||$q_{10}$|||||
| | 32 | 68 ||||||||||||||||||||||||||||||||$q_{10}$||||
| | 33 | 69 |||||||||||||||||||||||||||||||||$q_{10}$|||
| | 34 | 70 ||||||||||||||||||||||||||||||||||$q_{10}$||
| | 35 | 71 |||||||||||||||||||||||||||||||||||$q_{10}$|
| | 35+ | 72 ||||||||||||||||||||||||||||||||||||$q_{10}$|

| | | | State D | | | | | | | | | | | | | | | | | | | | | | | | | | | | | | | | | | | |
|---|---|---|---|---|---|---|---|---|---|---|---|---|---|---|---|---|---|---|---|---|---|---|---|---|---|---|---|---|---|---|---|---|---|---|---|---|---|---|
| | | n | 1 | 2 | 3 | 4 | 5 | 6 | 7 | 8 | 9 | 10 | 11 | 12 | 13 | 14 | 15 | 16 | 17 | 18 | 19 | 20 | 21 | 22 | 23 | 24 | 25 | 26 | 27 | 28 | 29 | 30 | 31 | 32 | 33 | 34 | 35 | 35+ |
| | n | | 37 | 38 | 39 | 40 | 41 | 42 | 43 | 44 | 45 | 46 | 47 | 48 | 49 | 50 | 51 | 52 | 53 | 54 | 55 | 56 | 57 | 58 | 59 | 60 | 61 | 62 | 63 | 64 | 65 | 66 | 67 | 68 | 69 | 70 | 71 | 72 |
| State E | 1 | 1 ||||||||||||||||||||||||||||||||||||
| | 2 | 2 ||||||||||||||||||||||||||||||||||||
| | 3 | 3 ||||||||||||||||||||||||||||||||||||
| | 4 | 4 ||||||||||||||||||||||||||||||||||||
| | 5 | 5 ||||||||||||||||||||||||||||||||||||
| | 6 | 6 ||||||||||||||||||||||||||||||||||||
| | 7 | 7 ||||||||||||||||||||||||||||||||||||
| | 8 | 8 ||||||||||||||||||||||||||||||||||||
| | 9 | 9 ||||||||||||||||||||||||||||||||||||
| | 10 | 10 ||||||||||||||||||||||||||||||||||||
| | 11 | 11 ||||||||||||||||||||||||||||||||||||
| | 12 | 12 ||||||||||||||||||||||||||||||||||||
| | 13 | 13 ||||||||||||||||||||||||||||||||||||
| | 14 | 14 ||||||||||||||||||||||||||||||||||||
| | 15 | 15 ||||||||||||||||||||||||||||||||||||
| | 16 | 16 ||||||||||||||||||||||||||||||||||||
| | 17 | 17 ||||||||||||||||||||||||||||||||||||
| | 18 | 18 ||||||||||||||||||||||||||||||||||||
| | 19 | 19 ||||||||||||||||||||||||||||||||||||
| | 20 | 20 ||||||||||||||||||||||||||||||||||||
| | 21 | 21 ||||||||||||||||||||||||||||||||||||
| | 22 | 22 ||||||||||||||||||||||||||||||||||||
| | 23 | 23 ||||||||||||||||||||||||||||||||||||
| | 24 | 24 ||||||||||||||||||||||||||||||||||||
| | 25 | 25 ||||||||||||||||||||||||||||||||||||
| | 26 | 26 ||||||||||||||||||||||||||||||||||||
| | 27 | 27 ||||||||||||||||||||||||||||||||||||
| | 28 | 28 ||||||||||||||||||||||||||||||||||||
| | 29 | 29 ||||||||||||||||||||||||||||||||||||
| | 30 | 30 ||||||||||||||||||||||||||||||||||||
| | 31 | 31 ||||||||||||||||||||||||||||||||||||
| | 32 | 32 ||||||||||||||||||||||||||||||||||||
| | 33 | 33 ||||||||||||||||||||||||||||||||||||
| | 34 | 34 ||||||||||||||||||||||||||||||||||||
| | 35 | 35 ||||||||||||||||||||||||||||||||||||
| | 35+ | 36 ||||||||||||||||||||||||||||||||||||
| State D | 1 | 37 ||$\gamma_1+\rho_1$||||||||||||||||||||||||||||||||||
| | 2 | 38 |$\delta_1$||$\gamma_1$|$\rho_1$||||||||||||||||||||||||||||||||
| | 3 | 39 ||$\delta_1$||$\gamma_1$||$\rho_1$||||||||||||||||||||||||||||||
| | 4 | 40 |||$\delta_1$||$\gamma_1$|||$\rho_1$||||||||||||||||||||||||||||
| | 5 | 41 ||||$\delta_1$||$\gamma_1$||||$\rho_1$||||||||||||||||||||||||||
| | 6 | 42 |||||$\delta_1$||$\gamma_1$|||||$\rho_1$||||||||||||||||||||||||
| | 7 | 43 ||||||$\delta_1$||$\gamma_1$||||||$\rho_1$||||||||||||||||||||||
| | 8 | 44 |||||||$\delta_1$||$\gamma_1$|||||||$\rho_1$||||||||||||||||||||
| | 9 | 45 ||||||||$\delta_1$||$\gamma_1$||||||||$\rho_1$||||||||||||||||||
| | 10 | 46 |||||||||$\delta_1$||$\gamma_1$|||||||||$\rho_1$||||||||||||||||
| | 11 | 47 ||||||||||$\delta_1$||$\gamma_1$||||||||||$\rho_1$||||||||||||||
| | 12 | 48 |||||||||||$\delta_1$||$\gamma_1$|||||||||||$\rho_1$||||||||||||
| | 13 | 49 ||||||||||||$\delta_1$||$\gamma_1$||||||||||||$\rho_1$||||||||||
| | 14 | 50 |||||||||||||$\delta_1$||$\gamma_1$|||||||||||||$\rho_1$||||||||
| | 15 | 51 ||||||||||||||$\delta_1$||$\gamma_1$||||||||||||||$\rho_1$||||||
| | 16 | 52 |||||||||||||||$\delta_1$||$\gamma_1$|||||||||||||||$\rho_1$||||
| | 17 | 53 ||||||||||||||||$\delta_1$||$\gamma_1$||||||||||||||||$\rho_1$||
| | 18 | 54 |||||||||||||||||$\delta_1$||$\gamma_1$||||||||||||||||$\rho_1$|
| | 19 | 55 ||||||||||||||||||$\delta_1$||$\gamma_1$|||||||||||||||$\rho_1$|
| | 20 | 56 |||||||||||||||||||$\delta_1$||$\gamma_1$||||||||||||||$\rho_1$|
| | 21 | 57 ||||||||||||||||||||$\delta_1$||$\gamma_1$|||||||||||||$\rho_1$|
| | 22 | 58 |||||||||||||||||||||$\delta_1$||$\gamma_1$||||||||||||$\rho_1$|
| | 23 | 59 ||||||||||||||||||||||$\delta_1$||$\gamma_1$|||||||||||$\rho_1$|
| | 24 | 60 |||||||||||||||||||||||$\delta_1$||$\gamma_1$||||||||||$\rho_1$|
| | 25 | 61 ||||||||||||||||||||||||$\delta_1$||$\gamma_1$|||||||||$\rho_1$|
| | 26 | 62 |||||||||||||||||||||||||$\delta_1$||$\gamma_1$||||||||$\rho_1$|
| | 27 | 63 ||||||||||||||||||||||||||$\delta_1$||$\gamma_1$|||||||$\rho_1$|
| | 28 | 64 |||||||||||||||||||||||||||$\delta_1$||$\gamma_1$||||||$\rho_1$|
| | 29 | 65 ||||||||||||||||||||||||||||$\delta_1$||$\gamma_1$|||||$\rho_1$|
| | 30 | 66 |||||||||||||||||||||||||||||$\delta_1$||$\gamma_1$||||$\rho_1$|
| | 31 | 67 ||||||||||||||||||||||||||||||$\delta_1$||$\gamma_1$|||$\rho_1$|
| | 32 | 68 |||||||||||||||||||||||||||||||$\delta_1$||$\gamma_1$||$\rho_1$|
| | 33 | 69 ||||||||||||||||||||||||||||||||$\delta_1$||$\gamma_1$|$\rho_1$|
| | 34 | 70 |||||||||||||||||||||||||||||||||$\delta_1$||$\gamma_1$|$\rho_1$|
| | 35 | 71 ||||||||||||||||||||||||||||||||||$\delta_1$||$\gamma_1+\rho_1$|
| | 35+ | 72 |||||||||||||||||||||||||||||||||||$\delta_1$||
