## Supplementary figures and images for "Over two orders of magnitude difference in rate of single chromosome loss among sundew (*Drosera* L., Droseraceae) lineages"

### Supplemental Figure 2

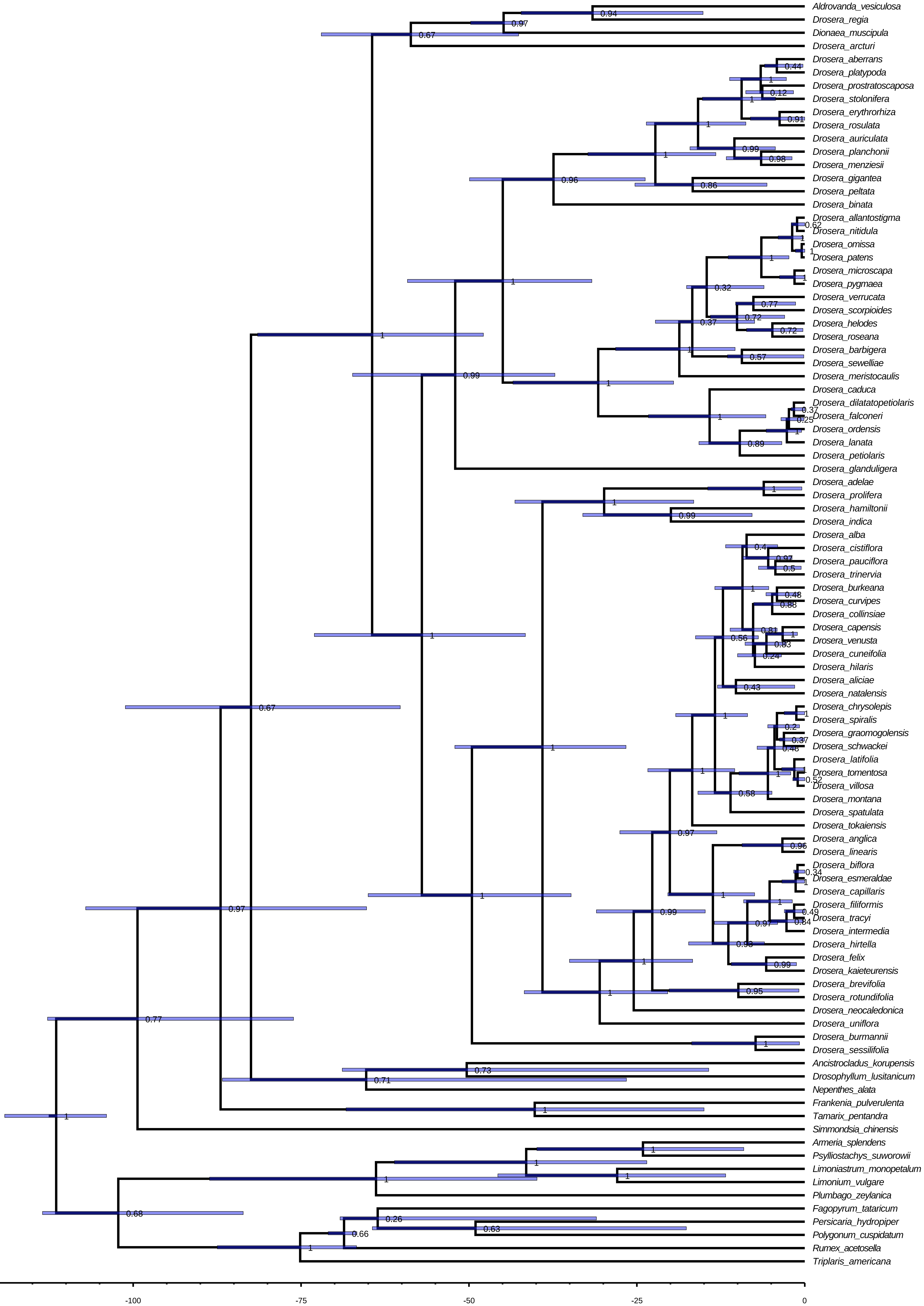
