## Supplemental Information 1 for "Over two orders of magnitude difference in rate of single chromosome loss among sundew (*Drosera* L., Droseraceae) lineages"

**Methods:**

As we were compiling chromosome counts, issues arose with species identification and chromosome count accuracy that were also concerns documented in other studies (Windham and Yatskievych 2003; Twardovska et al. 2008). Instead of simply removing chromosome counts with issues randomly (Goldblatt 2007), we developed a scoring system to quantify the degree of confidence and allow flexibility in the filtering criteria for downstream analyses. We used three scores to quantify three types of issues: a taxonomic score (confidence in species identification), chromosome count accuracy (confidence in count), and generalizability score (confidence in generalizing counts to a population). The scoring systems were tailored to meet our needs of analyzing genus level chromosome evolution by using only chromosome counts with high confidence scores.

Scoring followed the Supplemental Information S1 Table S1. The taxonomic score quantified our confidence of the correct species identification and ranged from 0 to 5. A low score indicates a lack of information or voucher, species misidentification, or changes in classification.

The chromosome count accuracy score ranged from 0 to 5. A low score indicates a low confidence of the correct chromosome count due to uncertainty in the exact number of chromosomes reported by the original authors, a few cells counted, and lack of an image documenting the source of the count (Windham and Yatskievych 2003).

Lastly, the generalizability score ranged from 0 to 4, and evaluated the ability to generalize observations as representative of a population. A low score indicates the count was from a hybrid, cultivated in tissue culture (Twardovska et al. 2008), or from a single plant. When multiple publications used the same cultivated source of a species or publications by the same author included the same species in multiple publications without a distinguishing location, the first publication was used, and all subsequent publications were treated as duplicates of the first publication.

Exceptions to the scoring matrix were documented in the Table S1 along with the full list of chromosome counts, scores, source of literature, voucher information, and notes.

| Supplemental Information S1 Table S1: The scoring guidelines used to evaluate confidence in score. Columns are the Taxonomic, Chromosome Count and Generalizability score. The rows are the different score values. Shaded cells were removed for analysis. | | | |
| --- | --- | --- | --- |
| Score | Taxonomic Score | Chromosome Count Accuracy Score | Generalizability Score Per Count Per Population |
| Ideal/Useful information | Voucher, collected from natural populations, provenance data, description of characters. | Number of cells counted, photos of chromosome squash. | Numbers of individuals counted per population, collected from wild. |
| 0 | NO species identified in original publication and no voucher for later identification OR confounding mismatch in species of voucher vs. chromosome count. | Uncertainty about chromosome counts in original publication. | (Natural OR artificial hybrids as unrepresentative of their taxon as each is a unique evolutionary event) OR Duplicate non-original counts already in the data set. |
| 1 | Species taxonomy changed since publication AND (NO location, OR character description, OR specimen to determine correct identity). | Only one cell was counted. | Clonally propagated individuals**. |
| 2 | (Species taxonomy changed since publication AND insufficient data for correct determination) OR species frequently mixed up/misidentified in cultivation | NO report of the number of cells counted AND NO image of squash. | (Cultivated OR Collected) AND (Counted 1 individual per population OR NO report of the number of individuals counted). |
| 3 | Species taxonomy is reportedly being reconsidered OR [a morphologically distinct species AND (NO location OR in cultivation)]. | (NO reports of the number of cells counted AND a photo of squashed cell*) OR 2 cells counted. | Collected AND Counted 2-4 individuals per population. |
| 4 | Voucher not verified but species uniquely characterized by location AND/OR a morphology (species unlikely to be confused with any other). | ((Photo of squashed cell*) AND 2 squashed cells) OR 3 cells counted. | Collected AND Counted >5 individuals per population. |
| 5 | Herbarium specimen verified by publication authors OR from the type author, specimen, or location. | 4+ cells counted. |  |
| *Photos of squashed cells have been reported as useful in correcting miscounts (Windham and Yatskievych 2003).  ** Clonal propagation has been reported to result in shifts in ploidy (Twardovska et al. 2008). | | | |
